# Spontaneous mutations conferring antibiotic resistance to antitubercular drugs at a range of concentrations in *Mycobacterium smegmatis*

**DOI:** 10.1101/296137

**Authors:** Iveren W Nyinoh, Johnjoe McFadden

## Abstract

Mycobacteria population can undergo mutations in their DNA sequence during replication, which if not repaired, would be transferred to future generations. In this study, *in vitro* spontaneous mutations in *Mycobacterium smegmatis* mc^2^155 (*Msm*) conferring resistance to isoniazid (INH^r^), rifampicin (RIF^r^), kanamycin (KAN^r^) and streptomycin (STR^r^) were determined at several concentrations in a fluctuation assay. Mutation rate was estimated using the *P₀* method, and estimates were then compared with the Lea-Coulson method of the median and Ma-Sandri-Sarkar Maximum Likelihood Estimator (MSS-MLE) method available on the Fluctuation analysis calculator (FALCOR). The mutation rates of RIF^r^ ranged from 9.24 × 10^-8^ - 2.18 × 10^-10^, INH^r^ 1.2 × 10^-7^ - 1.20×10^-9^, STR^r^ 2.77 × 10^-8^ - 5.31 × 10^-8^ and KAN^r^ 1.7 × 10^-8^ mutations per cell division. This study provides mutation rate estimates to key antitubercular drugs at a range of concentrations.

## Introduction

Antibiotic resistance allows bacteria to withstand the inhibitory effects of high concentrations of antibiotics, typically beyond the minimum concentration required to inhibit the growth of the bacterium. In mycobacteria, drug resistance occurs exclusively as a result of spontaneous mutations of genes encoding drug target (1), through a natural process of DNA replication. As such, resistant mutants can then be selected for in the presence of an antibiotic (2). *Mycobacterium tuberculosis* (*Mtb*) immune to several antitubercular drugs is referred to as multi-drug resistant and accounted for 490,000 cases of MDR-TB infections in 2017 and caused 190,000 deaths globally (3). The antibiotics used in this study are shown (Table 1, Figure 1).

**Table 1:**
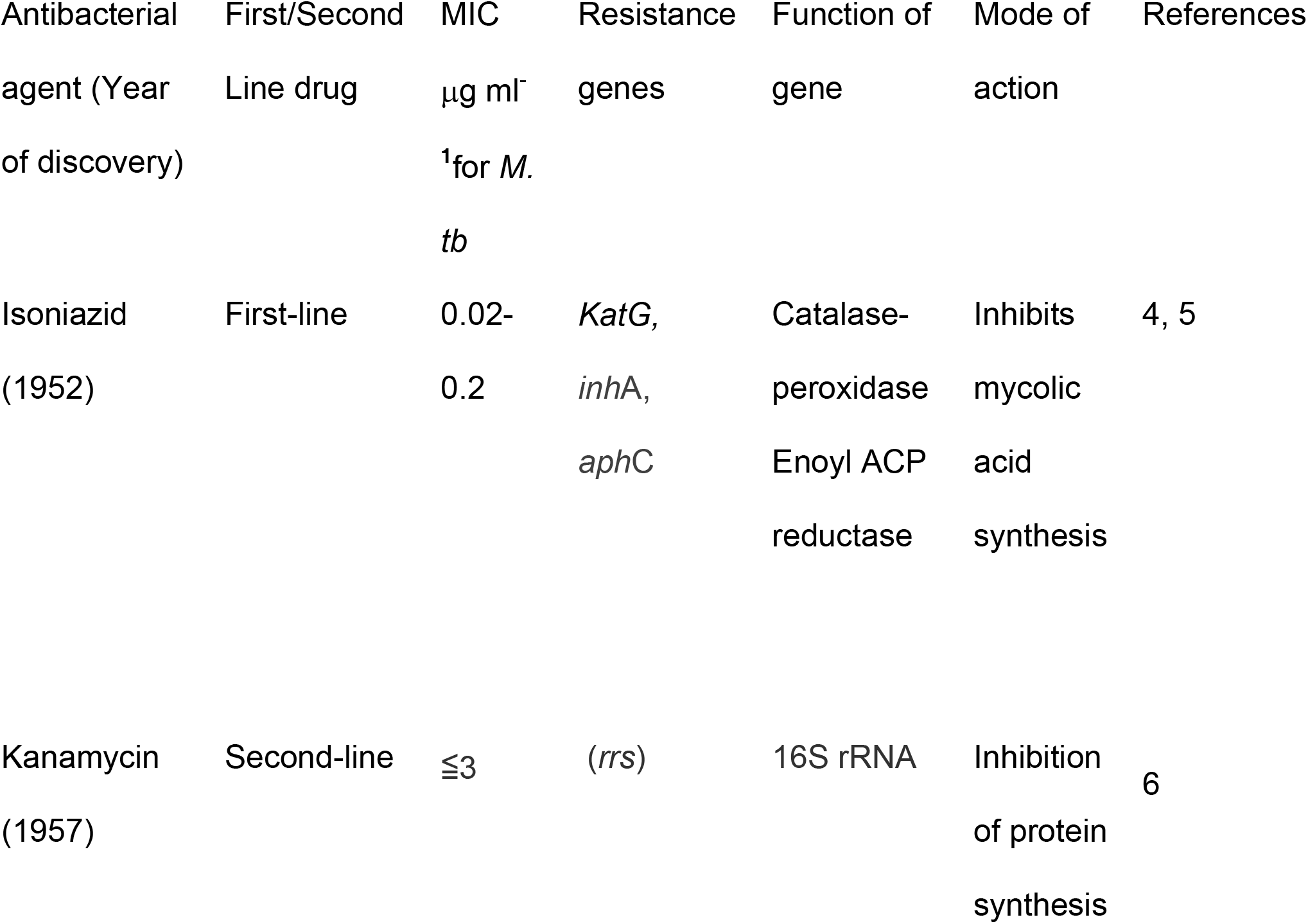
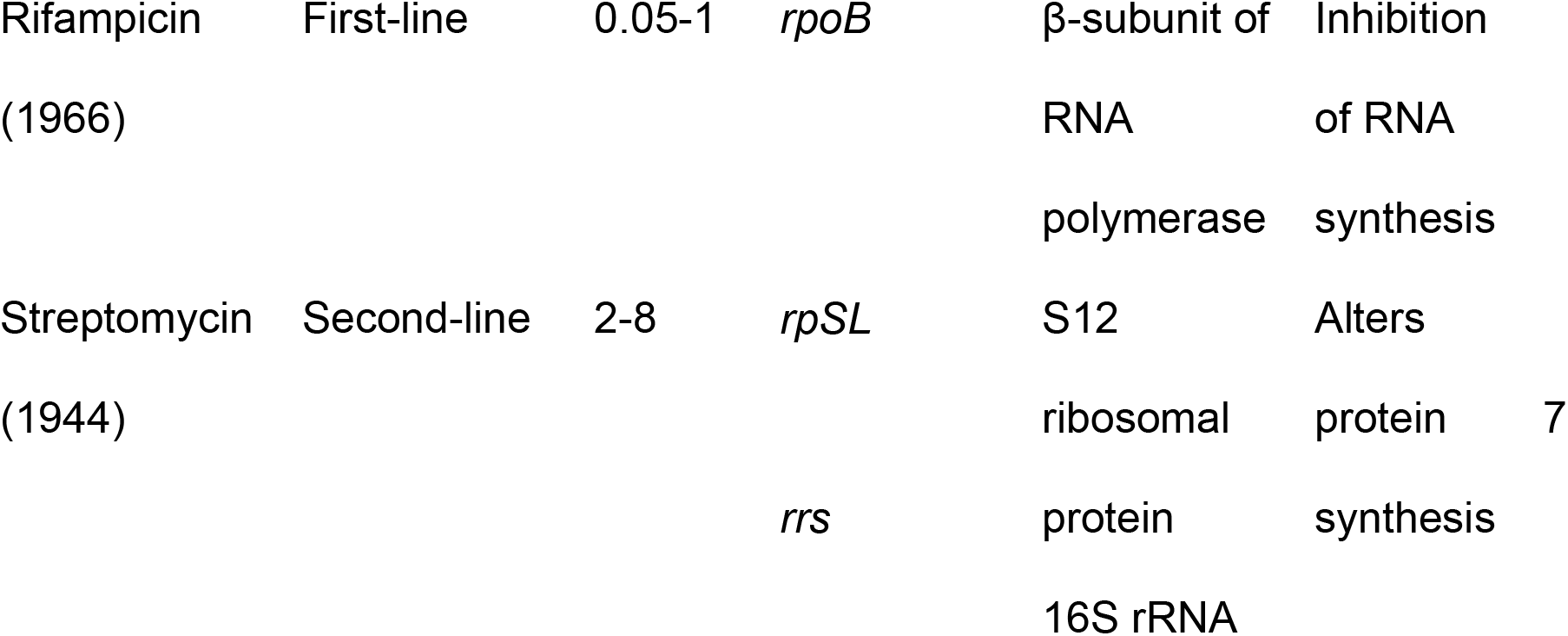
Modes of action of anti-TB drugs employed in this study.

**Figure 1:**
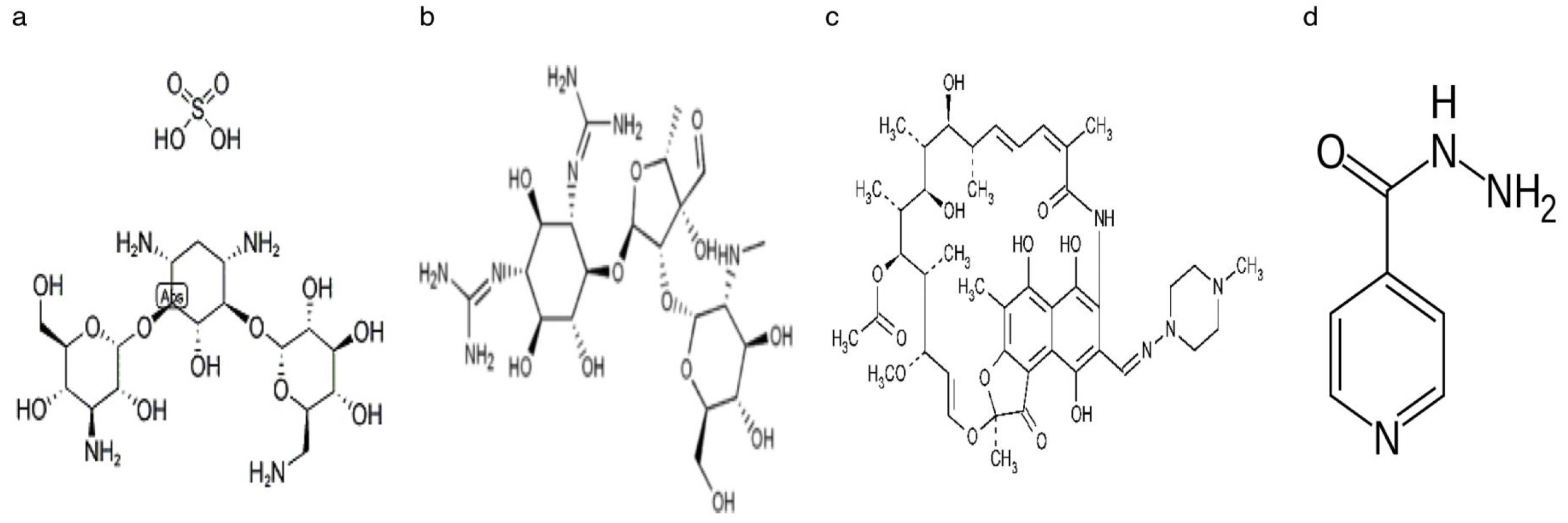
Structures of the anti-tubercular drugs used in this study-isoniazid (a) Kanamycin (b) Streptomycin (c) Rifampicin (d) Isoniazid (14).

Kanamycin (KAN) is an aminoglycoside antibiotic (Fig. 1a), belonging to the same oligosaccharide group of water-soluble antibiotic, as streptomycin (STR). Administered as a second-line anti TB drug, especially against STR-resistant *Mtb* (4), KAN is bactericidal and acts, by binding irreversibly to the 16s rRNA of the 30S ribosomal subunit thereby inhibiting the synthesis of protein (6). *In vitro* KAN-resistant *Mycobacterium smegmatis* (*Msm*) have been shown to possess altered ribosomes (9), which appear to be conferred by mutation in the *rrs* gene (6). STR (Fig. 1b) is an aminocyclitol aminoglycoside that inhibits translation during protein synthesis (6). Mutations in the *rpsL* gene and *rrs* genes result to STR-resistance (2, 10). RIF (Fig. 1c) binds to the β subunit of RNA polymerase, the enzyme responsible for transcription of RNA inhibiting transcription (11). RIF-resistance is due to altered β sub-unit of the polymerase at one of the three loci in *rpoB* gene (7). Together with RIF, INH (Fig. 1d) forms the core of TB therapy. Catalase-peroxidase enzyme (*katG*) activates INH into its active form and exerts its effect by inhibiting mycolic acid synthesis, which makes up a large amount of mycobacterial cell envelope (4). Generally, resistance to INH is often linked to a range of mutations affecting one or more genes e.g. mutations in the catalase-peroxidase gene, *KatG, inhA*-encoded long chain enoyl-ACP reductase, which is involved with the biosynthesis of mycolic acid (12). Recently also alkyl-hydroperoxide reductase (*aphC*) that is involved in the cellular response of oxidative stress (13, 5) has been reported to cause resistance in INH.

In this study, we isolated *Msm* mutants at a variety of concentrations and determined mutation rates using the *P₀* method (15), and estimates were then compared with the Lea-Coulson method of the median (LC-MM) and the Ma-Sandri-Sarkar Maximum Likelihood Estimator method (MSS-MLE), available on the fluctuation analysis calculator (16).

## Results

### Results for Minimum Inhibitory Concentration

Mutants are easier to detect at concentrations greater than the MIC, hence to find out what antibiotic concentration will result in the proliferation of resistant mutants, the minimum concentration, required to inhibit the visible growth of *Msm* was assayed. The results are shown (Table 2).

**Table 2:** Antitubercular drugs, targets and Minimum Inhibitory Concentrations results obtained in our study

| Drug | Symbol | Mechanism | MIC value |
| --- | --- | --- | --- |
| Isoniazid | INH | Mycolic Acid synthesis inhibitors | 8 $\mu\text{g ml}^{-1}$ |
| Kanamycin | KAN | Protein synthesis inhibitor | 0.24 $\mu\text{g ml}^{-1}$ |
| Rifampicin | RIF | DNA-dependent RNA synthesis inhibitor | 16 $\mu\text{g ml}^{-1}$ |
| Streptomycin | STR | Protein synthesis inhibitor | 0.5 $\mu\text{g ml}^{-1}$ |

### Results for fluctuation assay

Table 3 shows the results for mutation rate calculated using the *P₀* method.

**Table 3:** Results for mutation rate using the *P*_*0*_ method

| Mutant | Conc.<br>$\mu\text{g ml}^{-1}$ | Mean | Cultures | Zeros | Variance | SD | No. of<br>viable<br>bacteria<br>($Nt$) | $P_0$ | $m$ | Mutation<br>rate |
| --- | --- | --- | --- | --- | --- | --- | --- | --- | --- | --- |
| RIF<sup>r</sup> | 100 | 20.5 | 10 | 03 | 2323.39 | 48.20 | 9.00E+06 | 0.3 | 1.20 | $9.24 \times 10^{-8}$ |
| RIF<sup>r</sup> | 200 | 8.5 | 24 | 08 | 247.22 | 15.72 | 1.72E+09 | 0.33 | 1.11 | $4.47 \times 10^{-10}$ |
| RIF<sup>r</sup> | 500 | 1.42 | 24 | 14 | 8.60 | 2.93 | 1.72E+09 | 0.58 | 0.54 | $2.18 \times 10^{-10}$ |
| INH<sup>r</sup> | 500 | 30.4 | 10 | 02 | 1901.6 | 43.61 | 9.00E+06 | 0.2 | 1.61 | $1.24 \times 10^{-7}$ |
| INH<sup>r</sup> | 1000 | 7.9 | 20 | 01 | 94.2 | 9.71 | 1.72E+09 | 0.05* | 2.99 | $1.20 \times 10^{-9}$ |
| STR<sup>r</sup> | 20 | 0.5 | 10 | 07 | 0.72 | 0.85 | 9.00E+06 | 0.7 | 0.36 | $2.77 \times 10^{-8}$ |
| STR<sup>r</sup> | 100 | 0.8 | 10 | 05 | 0.84 | 0.92 | 9.00E+06 | 0.5 | 0.69 | $5.31 \times 10^{-8}$ |
| KAN<sup>r</sup> | 100 | 0.2 | 10 | 08 | 0.18 | 0.42 | 9.00E+06 | 0.8* | 0.22 | $1.70 \times 10^{-8}$ |

Mutant = antibiotic used in mutant selection; <sup>r</sup> = resistance; Conc. $\mu\text{g/ml}$ represents the concentrations used in the selection of mutants; Mean= average of the number of mutants in the cultures. The mean was calculated by dividing the total number of colonies by the number of cultures; Cultures = number of parallel independent cultures used in the study for each antibiotic at the defined concentration; Zeros = number of cultures with zero or no mutants; Variance = the extent to which the individual numbers of mutants are offset from the mean; Standard deviation ( $SD$ ) is calculated as the square root of the variance and
shows how much the number of mutants deviates from the mean; No. of bacteria ( $N_t$ ) = final number of cells plated; $P_0$ = Proportion of cultures without mutants (obtained by dividing the zeros by the number of cultures); Values in \* are those not within the accepted values for the $P_0$ method ( $0.7 \geq P_0 \geq 0.1$ ( $0.3$ $\leq m \leq 2.3$ ); $m$ = number of mutations per culture, obtained by taking the negative natural log of $P_0$ ; Mutation rate calculated is per cell per generation using $m$ $\times \ln 2 / N_t$ .

### Results for mutation rate using Fluctuation Analysis Calculator (FALCOR)

Due to the fact that the *P*_*0*_ method is not applicable across all values of *m* i.e. the number of mutations per culture (15), the LC-MM and MSS-MLE, available on the Fluctuation analysis calculator (16) were also employed in estimating mutation rate of *Msm* to the test antibiotics (Table 4) and then and compared with the *P*_*0*_ method.

**Table 4:**
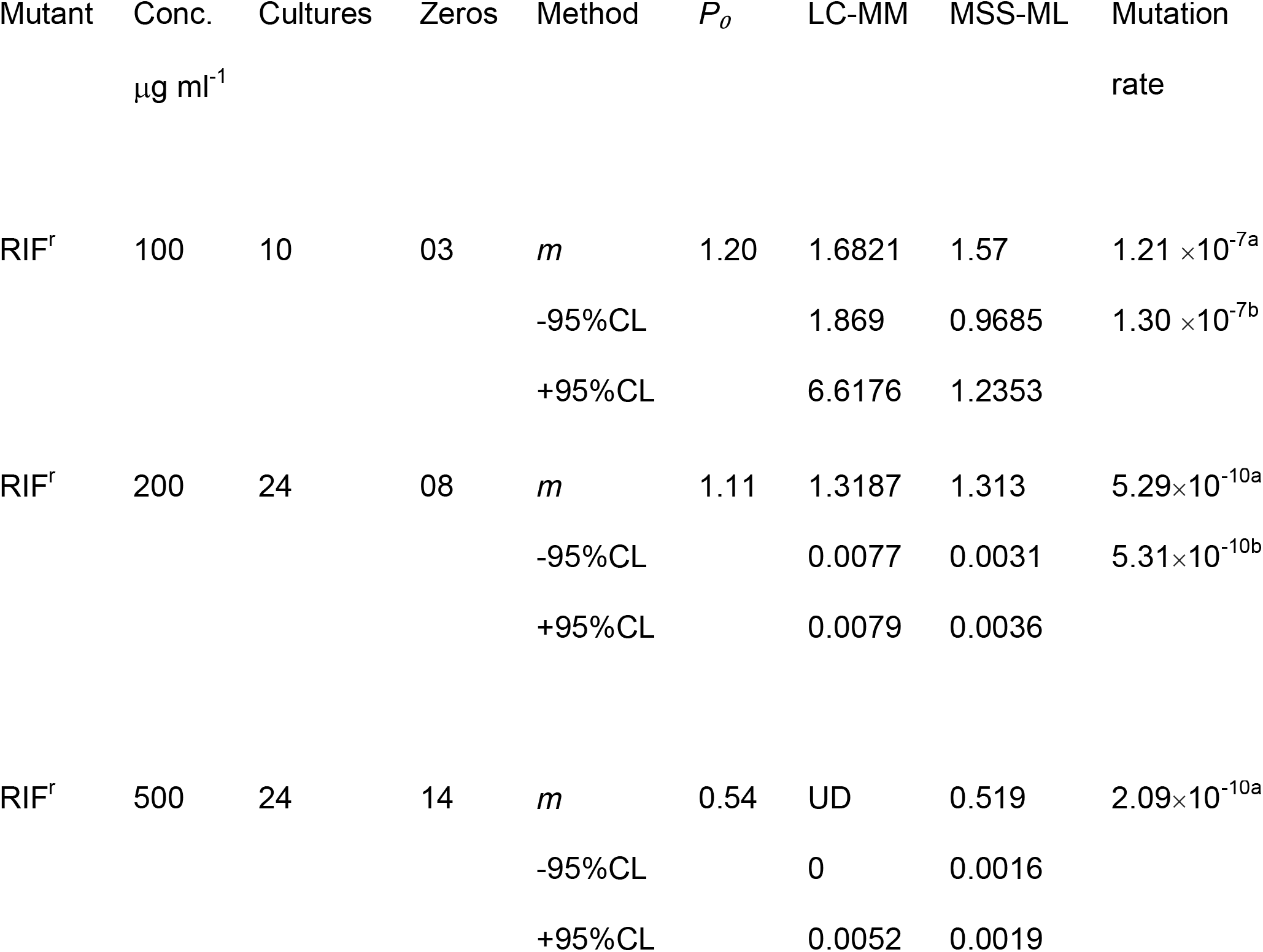
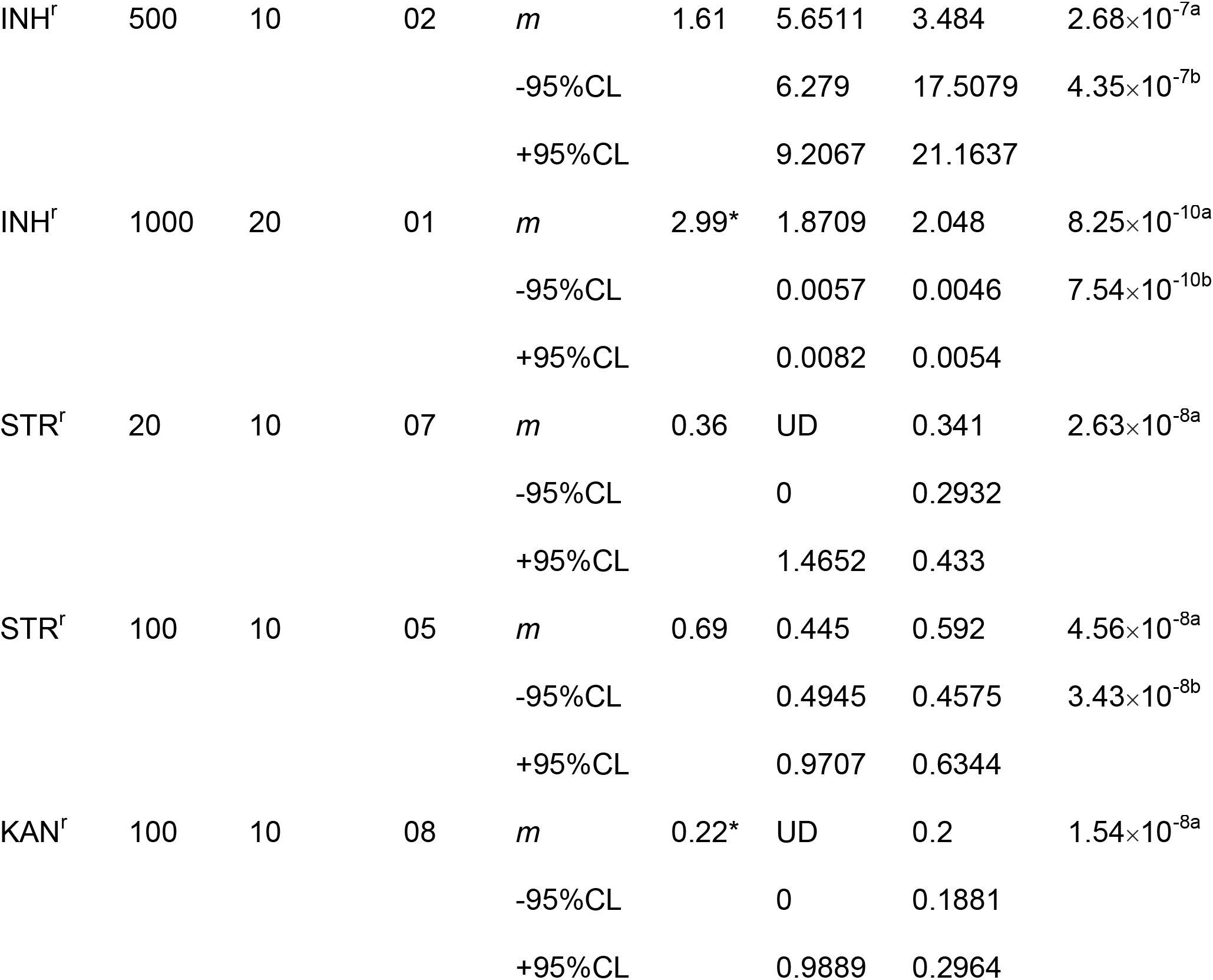
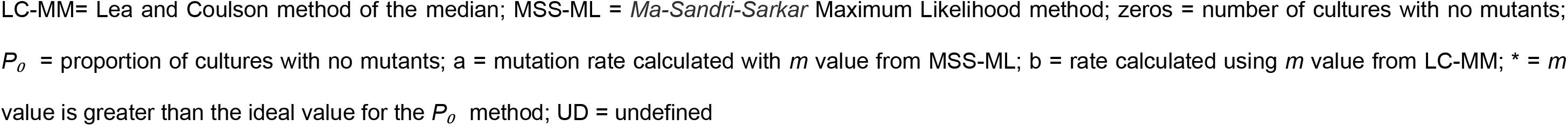
Evaluation of *P*_*0*_ method against two other methods available on the Fluctuation analysis calculator with experimental data

| Mutant | Conc.<br>$\mu\text{g ml}^{-1}$ | Cultures | Zeros | Method | $P_0$ | LC-MM | MSS-ML | Mutation<br>rate |
| --- | --- | --- | --- | --- | --- | --- | --- | --- |
| RIF<sup>r</sup> | 100 | 10 | 03 | $m$ | 1.20 | 1.6821 | 1.57 | $1.21 \times 10^{-7a}$ |
| | | | | -95%CL | | 1.869 | 0.9685 | $1.30 \times 10^{-7b}$ |
|  |  |  |  | +95%CL |  | 6.6176 | 1.2353 |  |
| RIF<sup>r</sup> | 200 | 24 | 08 | $m$ | 1.11 | 1.3187 | 1.313 | $5.29 \times 10^{-10a}$ |
| | | | | -95%CL | | 0.0077 | 0.0031 | $5.31 \times 10^{-10b}$ |
|  |  |  |  | +95%CL |  | 0.0079 | 0.0036 |  |
| RIF<sup>r</sup> | 500 | 24 | 14 | $m$ | 0.54 | UD | 0.519 | $2.09 \times 10^{-10a}$ |
|  |  |  |  | -95%CL |  | 0 | 0.0016 |  |
|  |  |  |  | +95%CL |  | 0.0052 | 0.0019 |  |
| INH<sup>r</sup> | 500 | 10 | 02 | *m* | 1.61 | 5.6511 | 3.484 | 2.68×10<sup>-7a</sup> |
|  |  |  |  | -95%CL |  | 6.279 | 17.5079 | 4.35×10<sup>-7b</sup> |
|  |  |  |  | +95%CL |  | 9.2067 | 21.1637 |  |
| INH<sup>r</sup> | 1000 | 20 | 01 | *m* | 2.99* | 1.8709 | 2.048 | 8.25×10<sup>-10a</sup> |
|  |  |  |  | -95%CL |  | 0.0057 | 0.0046 | 7.54×10<sup>-10b</sup> |
|  |  |  |  | +95%CL |  | 0.0082 | 0.0054 |  |
| STR<sup>r</sup> | 20 | 10 | 07 | *m* | 0.36 | UD | 0.341 | 2.63×10<sup>-8a</sup> |
|  |  |  |  | -95%CL |  | 0 | 0.2932 |  |
|  |  |  |  | +95%CL |  | 1.4652 | 0.433 |  |
| STR<sup>r</sup> | 100 | 10 | 05 | *m* | 0.69 | 0.445 | 0.592 | 4.56×10<sup>-8a</sup> |
|  |  |  |  | -95%CL |  | 0.4945 | 0.4575 | 3.43×10<sup>-8b</sup> |
|  |  |  |  | +95%CL |  | 0.9707 | 0.6344 |  |
| KAN<sup>r</sup> | 100 | 10 | 08 | *m* | 0.22* | UD | 0.2 | 1.54×10<sup>-8a</sup> |
|  |  |  |  | -95%CL |  | 0 | 0.1881 |  |
|  |  |  |  | +95%CL |  | 0.9889 | 0.2964 |  |

LC-MM= Lea and Coulson method of the median; MSS-ML = *Ma-Sandri-Sarkar* Maximum Likelihood method; zeros = number of cultures with no mutants; $P_0$ = proportion of cultures with no mutants; a = mutation rate calculated with $m$ value from MSS-ML; b = rate calculated using $m$ value from LC-MM; \* = $m$ value is greater than the ideal value for the $P_0$ method; UD = undefined

## Discussion

We have isolated mutants to *Msm* at a range of antitubercular drug concentrations. RIF was tested at 100, 200 and 500 µg ml^-1^, INH at 500 and 1000 µg ml^-1^, STR at 20 and 100 µg ml^-1^ and KAN at 100 µg ml^-1^ using the fluctuation assay. We next estimated mutation rates using the *P*_*0*_ method, and estimates were then compared with LC-MM and MSS-MLE methods obtainable on the fluctuation analsysis calculator. In our study, the mutation rates estimated using the three methods were comparable. For instance at 100 µg ml^-1^ KAN, the mutation rate was 1.70 × 10^-8^ mutations per cell division with the *P*_*0*_ method and 1.54 10^-8^ with the LC-MM.

The hypothesis surrounding spontaneous mutation predicts a large fluctuation around the average for the count taken from the individual cultures. As such, a mutation occurring earlier in the growth of the culture results in a higher number of mutated cells (17). This was observed for 500 µg ml^-1^ INH and mutation rate in our study was significantly raised (10^-7^) for post log phase growth. Complex networks of factors influence the rate and type of mutants that can be selected with a given antibiotic. One of such factors that play a significant part in the mutation rate is the concentration of the antibiotic (18, 19). Thus it could be observed that, when the concentration of RIF increased from 100 to 200 and 500 µg ml^-1^ on agar plates, the number of mutants selected reduced and the rate of mutations ranged from 9.24 × 10^-8^, 4.47 × 10^-10^, and 2.18 ×10^-10^. Respectively for estimation using the *P*_*0*_ method, LC-MM and MSS-MLE.

In a study by (20), mutation frequencies of *Msm* mc^2^6, rather than mutation rate was estimated, to 100 µg ml^-1^ STR and 500 µg ml^-1^ RIF and were >2 × 10^-4^ and >2.4 ×10^-5^ respectively which were higher than the results obtained in the current study. However, the *Msm* strains employed in both studies were different (mc^2^6 versus mc^2^155). It was difficult to obtain STR^r^ and KAN^r^ mutants. Causes of resistance in streptomycin have been extensively investigated in many bacteria and require a very specific base substitution in ribosomal genes for one of the ribosomal proteins. Mutations in the 16srRNA gene *rrs* have been found to confer mutation in streptomycin and kanamycin (6). Hence the mutation rates calculated using the *P_o_* method for this study were low i.e. 2.77 × 10^-8^ for streptomycin at 20 μg ml^1^ and 5.31 × 10^-8^ for streptomycin at 100 μg ml^-1^. These results are comparable to streptomycin mutation rate of 2.29 × 10^-8^ in *Mtb* (2). Mutation rate of *Msm* to 100 μg ml^-1^ KAN was 1.70 ×10^-8^ in this study. Just as it occurs in *Mtb*, rifampicin resistance mutation rate of 3.32 × 10^-9^ (2) has been found to be similar to *Msm* mutation rate of 10^-8^-10^-10^ as we have found in our study.

## Conclusions

It was not surprising that *M. smegmatis* been naturally resistant to isoniazid had the lowest mutation rate by comparison to kanamycin, streptomycin and rifampicin. Furthermore, we found that since the L-C method estimates mutation rate based on the median value, it was not ideal for estimating mutation rate where greater than 50 per cent of cultures had no growth (15). This resulted in both an undefined *m* value and uncalculated mutation rate. All the three methods were ideal in estimating mutation rate provided the recommendations for *P*_*0*_ and *m* values are adhered to. These findings will promote a greater understanding of the phenomenon of mutation and how the estimation of mutation rate could be of importance in the control of multi-drug resistant *M. tuberculosis* infection.

## Materials and Methods Bacteria strain

*Mycobacterium smegmatis* mc^2^155 was employed in our experiments (21).

### Test drugs

Isoniazid, kanamycin, Rifampicin and Streptomycin all purchased from Sigma-Aldrich Chemical Co. Ltd. (Poole, UK) were employed in this study. Standard stock solutions of isoniazid (50 mg/mL), Kanamycin (10 mg/mL) and Streptomycin (10mg/mL) were prepared by dissolving in sterile distilled water (SDW) and filtering using a 0.22 micrometre (µm) pore size cellulose membrane, while RIF (50 mg/mL) stock solution was prepared in DMSO (Fisher Scientific Ltd. Leicestershire, UK). Working solutions were prepared by diluting in SDW.

### Growth medium

Nutrient broth No. 2 (NB2; Lab lemco powder 10, Oxoid Ltd, Basingstoke, England), composed of: peptone 10g, sodium chloride 5g, beef extract 5g and reverse osmosis water, to make 1000 mL. Agar Technical No. 3 (Oxoid Ltd, Basingstoke, England) at 1.5 % was used. Sterilisation was by autoclaving at 121°C, 15 pound per square inch (psi) for 20 minutes. Antibiotics were added to media after cooling to 55°C.

### Supplements

Glycerol 0.5 % volume by volume (v/v), 0.1 % (v/v) Tween 80 was used to supplement the broth. DMSO was also used in rifampicin stock solution preparation (Fisher Scientific Ltd, Leicestershire, UK).

### Determination of Minimum Inhibitory Concentration (MIC)

MIC assays were determined based on log_2_ serial dilution of broth using NB2 containing 0.1 % Tween 80 in 5 mL tubes using the procedure by (22). This was incubated for 48 h at 37°C and tubes were then observed for visible growth.

### Fluctuation assay

The distribution of mutant numbers in parallel cultures was determined using Fluctuation analysis method. A small number of cells (OD, 0.002) were grown under non-selective conditions in a 15 mL centrifuge tube. After about 36 h, a 1:1 serial dilution was done appropriately by introducing 1 mL of inoculum into 1 mL sterile NB2 medium. This was to ensure the numbers of cells in all tubes were the same. About 10 to 24 parallel independent bijou bottles containing 2 mL were used in the fluctuation assay. Thereafter, the cells were grown to saturation (after 4 to 7 days), resulting in 10^6^ to 10^9^ cells, and selected for mutant growth. Microfuge tubes (1.5 mL) were then used to pellet the culture (10 000 g, 5 minutes) and made into 300 L volume. 200 µL volume was plated on antibiotic selective plates at the indicated concentrations. In order to estimate the number of viable cells, the remaining 100 µL was serially diluted and plated on non-selective NB2 agar plates. The average cell number was then calculated. This was the final number of cells, *Nt* that was used for mutation rate calculation.

### Isolation of antibiotic-resistant mutants

*Msm* mc^2^155 cultures were spread out on NBA plates at the desired antibiotic concentrations and incubated for 4-6 days at 37 °C. Mutant colonies were confirmed by re-streaking on antibiotic selective plates containing the antibiotics at their respective concentrations.

## Data analysis

The numbers of mutant cells were analysed using *P*_*0*_ method and fluctuation analysis calculator (FALCOR) available online.

